# Extracting a Biologically Relevant Latent Space from Cancer Transcriptomes with Variational Autoencoders

**DOI:** 10.1101/174474

**Authors:** Gregory P. Way, Casey S. Greene

## Abstract

The Cancer Genome Atlas (TCGA) has profiled over 10,000 tumors across 33 different cancer-types for many genomic features, including gene expression levels. Gene expression measurements capture substantial information about the state of each tumor. Certain classes of deep neural network models are capable of learning a meaningful latent space. Such a latent space could be used to explore and generate hypothetical gene expression profiles under various types of molecular and genetic perturbation. For example, one might wish to use such a model to predict a tumor’s response to specific therapies or to characterize complex gene expression activations existing in differential proportions in different tumors. Variational autoencoders (VAEs) are a deep neural network approach capable of generating meaningful latent spaces for image and text data. In this work, we sought to determine the extent to which a VAE can be trained to model cancer gene expression, and whether or not such a VAE would capture biologically-relevant features. In the following report, we introduce a VAE trained on TCGA pan-cancer RNA-seq data, identify specific patterns in the VAE encoded features, and discuss potential merits of the approach. We name our method “Tybalt” after an instigative, cat-like character who sets a cascading chain of events in motion in Shakespeare’s “*Romeo and Juliet*”. From a systems biology perspective, Tybalt could one day aid in cancer stratification or predict specific activated expression patterns that would result from genetic changes or treatment effects.

## 1. Introduction

Deep learning has improved the state of the art in many domains, including image, speech, and text processing, but it has yet to make significant enough strides in biomedicine for it to be considered transformative.^1^ Nevertheless, several studies have revealed promising results. For instance, Esteva *et al.* used convolutional neural networks (CNNs) to diagnose melanoma from skin images and Zhou and Troyanskaya trained deep models to predict the impact of noncoding variants.^2,3^ However, several domain specific limitations remain. In contrast to image or text data, validating and visualizing learning in biological datasets is particularly challenging. There is also a lack of ground truth labels in biomedical domains, which often limits the efficacy of supervised models. New unsupervised deep learning approaches such as generative adversarial nets (GANs) and variational autoencoders (VAEs) harness the modeling power of deep learning without the need for accurate labels.^4^–^6^ Unlike traditional CNNs, which model data by minimizing inaccurate class predictions, autoencoder models, including VAEs, learn through data reconstruction. Reconstructing gene expression input data using autoencoder frameworks has been previously shown to reveal novel biological patterns.^7^–^9^

VAEs and GANs are generative models, which means they learn to approximate a data generating distribution. Through approximation and compression, the models have been shown to capture an underlying data manifold — a constrained, lower dimensional space where data is distributed — and disentangle sources of variation from different classes of data.^10,11^ For instance, a recent group trained adversarial autoencoders on chemical compound structures and their growth inhibiting effects in cancer cell lines to learn manifold spaces of effective small molecule drugs.^12,13^ Additionally, Rampasek *et al.* trained a VAE to learn a gene expression manifold of reactions of cancer cell lines to drug treatment perturbation.^14^ The theoretical basis for modeling cancer using lower dimensional manifolds is established, as it has been previously hypothesized that cancer exists in “basins of attraction” defined by specific pathway aberrations that drive cells toward cancer states.^15^ These states could be revealed by data driven manifold learning approaches.

The Cancer Genome Atlas (TCGA) has captured several genomic measurements for over 10,000 different tumors across 33 cancer-types.^16^ TCGA has released this data publicly, enabling many secondary analyses, including the training of deep models that predict survival.^17^ One data type amenable to modeling manifold spaces is RNA-seq gene expression because it can be used as a proxy to describe tumor states and the downstream consequences of specific molecular aberration. Biology is complex, consisting of multiple nonlinear and often redundant connections among genes, and when a specific pathway aberration occurs, the downstream response to the perturbation is captured in the transcriptome. In the following report, we extend the autoencoder framework by training and evaluating a VAE on TCGA RNA-seq data. We aim to demonstrate the validity and specific latent space benefits of a VAE trained on gene expression data. We do not aim to comprehensively profile all learned pan-cancer VAE features nor survey clinical implications. We also do not compare our approach to alternate dimensionality reduction algorithms, but instead present our model as an additional tool in the toolkit for extracting knowledge from gene expression. We shall name this model “Tybalt”.

## 2. Methods

### 2.1 Model Summary

VAEs are data driven, unsupervised models that can learn meaningful latent spaces in many contexts. In this work, we aim to build a VAE that compresses gene expression features and reveals a biologically relevant latent space. The VAE is based on an autoencoding framework, which can discover nonlinear explanatory features through data compression and nonlinear activation functions. A traditional autoencoder consists of an encoding phase and a decoding phase where input data is projected into lower dimensions and then reconstructed.^18^ An autoencoder is deterministic, and is trained by minimizing reconstruction error. In contrast, VAEs are stochastic and learn the *distribution* of explanatory features over samples. VAEs achieve these properties by learning two distinct latent representations: a mean and standard deviation vector encoding. The model adds a Kullback-Leibler (KL) divergence term to the reconstruction loss, which also regularizes weights through constraining the latent vectors to match a Gaussian distribution. In a VAE, these two representations are learned concurrently through the use of a reparameterization trick that permits a back propagated gradient.^4^ Importantly, new data can be projected onto an existing VAE feature space enabling new data to be assessed.

### 2.2 Model Implementation

VAEs have been shown to generate “blurry” data compared with other generative models, including GANs, but VAEs are also generally more stable to train.^19^ We trained our VAE model, Tybalt, with the following architecture: 5,000 input genes encoded to 100 features and reconstructed back to the original 5,000 (Figure 1A). The 5,000 input genes were selected based on highest variability by median absolute deviation (MAD) in the TCGA pan-cancer dataset.

We initially trained Tybalt without batch normalization,^20^ but observed that when we included batch normalization in the encoding step, we trained faster and with heterogeneous feature activation. Batch normalization in machine learning is distinct from normalizing gene expression batches together in data processing. In machine learning, batch normalization adds additional feature regularization by scaling activations to zero mean and unit variance, which has been observed to speed up training and reduce batch to batch variability thus increasing generalizability. We trained Tybalt with an Adam optimizer,^21^ included rectified linear units^22^ and batch normalization in the encoding stage, and sigmoid activation in the decoding stage. We built Tybalt in Keras (version 2.0.6)^23^ with a TensorFlow backend (version 1.0.1).^24^ For more specific VAE illustrations and walkthroughs refer to an extended tutorial^25^ and these intuitive blog posts.^26,27^

### 2.3 Parameter Selection

We performed a parameter sweep over batch size (50, 100, 128, 200), epochs (10, 25, 50, 100), learning rates (0.005, 0.001, 0.0015, 0.002, 0.0025) and warmups (*κ*) (0.01, 0.05, 0.1, and 1). *κ* controls how much the KL divergence loss contributes to learning, which effectively transitions a deterministic autoencoder to a VAE.^28,29^ For instance, a *κ* = 0.1 would add 0.1 to a weight on the KL loss after each epoch. After 10 epochs, the KL loss will have equal weight as the reconstruction loss. We did not observe *κ* to influence model training (Figure 1B), so we kept *κ* = 1 for downstream analyses. We evaluated train and test set loss at each epoch. The test set was a random 10% partition of the full data. In general, training was relatively stable for many parameter combinations, but was consistently worse for larger batches, particularly with low learning rates. Ultimately, the best parameter combination based on validation loss was batch size 50, learning rate 0.0005, and 100 epochs (Figure 1C). Because training stabilized after about 50 epochs, we terminated training early. Training and testing loss across all 50 epochs is shown in Figure 1D. We performed the parameter sweep on a cluster of 8 NVIDIA GeForce GTX 1080 Ti GPUs on the PMACS cluster at The University of Pennsylvania.

**Fig. 1.**
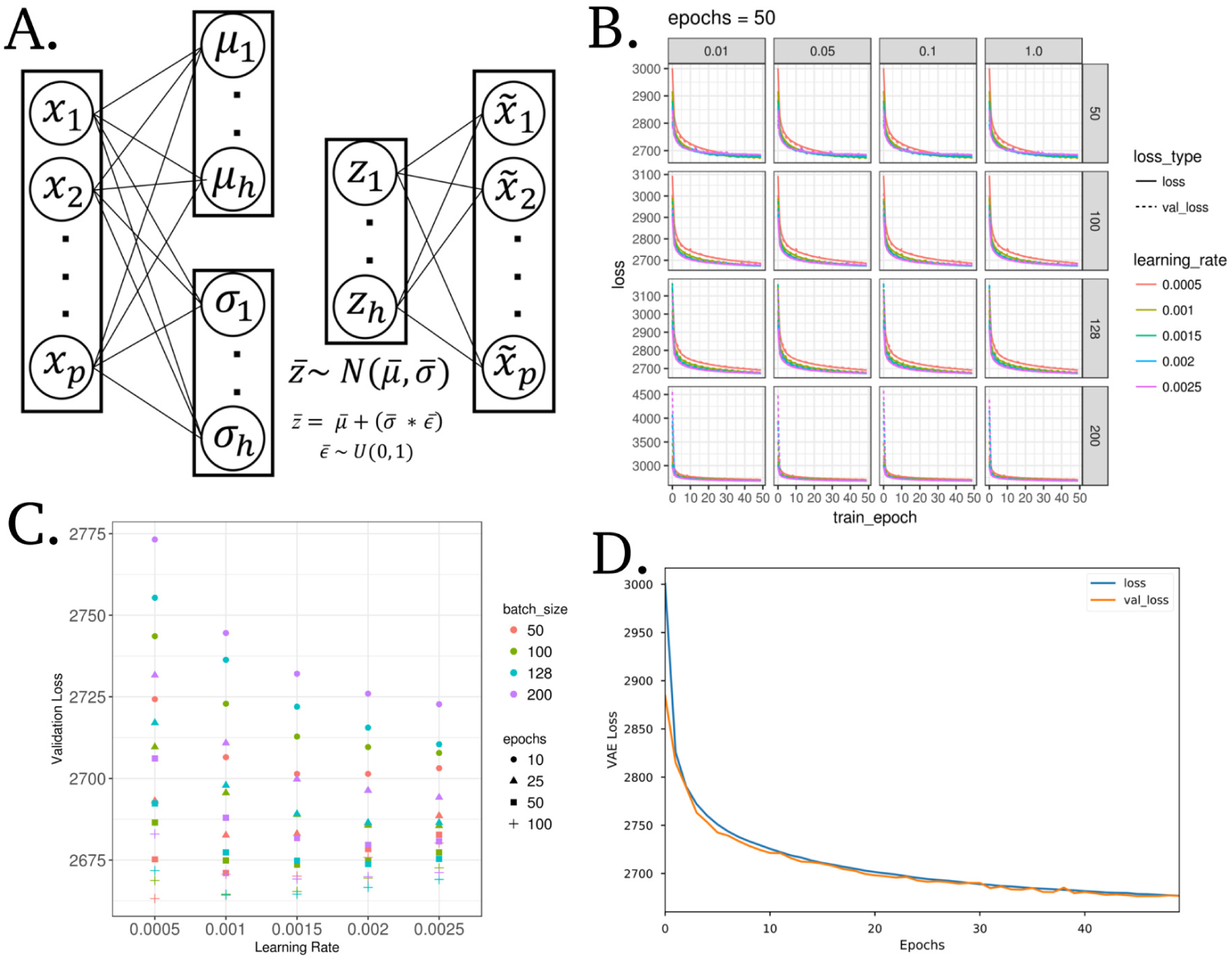
A variational autoencoder (VAE) applied to model gene expression data. **(A)** Model wire diagram of Tybalt encoding a gene expression vector (*p* = 5,000) into mean (*μ*) and standard deviation (*σ*) vectors (*h* = 100). A reparameterization trick^4,5^ enables learning *z*, which is then reconstructed back to input 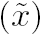. **(B)** Training and validation VAE loss across training epochs (full pass through all training data). Shown across vertical and horizontal facets are values of *κ* and batch size, respectively. **(C)** Final validation loss for all parameters with *κ* = 1. **(D)** VAE loss for training and testing sets through optimized model training.

### 2.4 Input Data

The input data consisted of level 3 TCGA RNA-seq gene expression data for 9,732 tumors and 727 tumor adjacent normal samples (10,459 total samples) measured by the 5,000 most variably expressed genes. The full dataset together is referred to as the pan-cancer data. The level 3 RNA-seq data consists of a preprocessed and batch-corrected gene abundance by sample matrix measured by log2(FPKM + 1) transformed RSEM values. The most variably expressed genes were defined by median absolute deviation (MAD). In total, there were 33 different cancer-types (including glioblastoma, ovarian, breast, lung, bladder cancer, etc.) profiled, each with varying number of tumors. We accessed RNA-seq data from the UCSC Xena data browser on March 8th, 2016 and archived the data in Zenodo.^30^ To facilitate training, we min-maxed scaled RNA-seq data to the range of 0 - 1. We used corresponding clinical data accessed from the Snaptron web server.^31^

### 2.5 Interpretation of Gene Weights

Much like the weights of a deterministic autoencoder, Tybalt’s decoder weights captured the contribution of specific genes to each learned feature.^7,8,32^ For most features, the distribution of gene weights was similar: Many genes had weights near zero and few genes had high weights at each tail. In order to characterize patterns explained by selected encoded features of interest, we performed overrepresentation pathway analyses (ORA) separately for both positive and negative high weight genes; defined by greater than 2.5 standard deviations above or below the mean, respectively. We used WebGestalt,^33^ with a background of the 5,000 assayed genes, to perform the analysis over gene ontology (GO) biological process terms.^34^ P values are presented after an Benjamini-Hochberg FDR adjustment.

### 2.6 The Latent Space of Ovarian Cancer Subtypes

Image processing studies have shown the remarkable ability of generative models to mathematically manipulate learned latent dimensions.^35,36^ For example, subtracting the image latent representation of a neutral man from a smiling man and adding it to a neutral woman, resulted in a vector associated with a smiling woman. We were interested in the extent to which Tybalt learned a manifold representation that could be manipulated mathematically to identify state transitions across high grade serous ovarian cancer (HGSC) subtypes. The TCGA naming convention of these subtypes is mesenchymal, proliferative, immunoreactive, and differentiated.^37^

To characterize the largest differences between the mesenchymal/immunoreactive and proliferative/differentiated HGSC subtypes, we performed a series of mean HGSC subtype vector subtractions in Tybalt latent space:

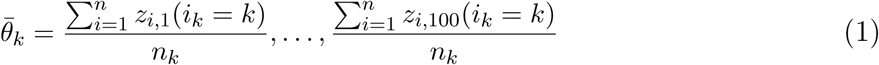

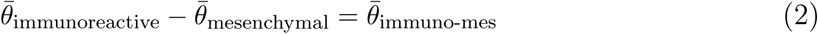

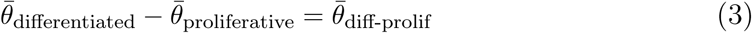

Where (*i*_*k*_ = *k*) is an indicator function if sample *i* has membership with subtype *k* and *z* is the encoded layer. We used tumor subtype assignments provided for TCGA samples in Verhaak *et al.* 2013.^38^ If Tybalt learned a biological manifold, this subtraction would result in the identification of biologically relevant features stratifying tumors of specific subtypes with a continuum of expression states.

### 2.7 Enabling Exploration through Visualization

We provide a Shiny app to interactively visualize activation patterns of encoded Tybalt features with covariate information at <u>https://gregway.shinyapps.io/pancan_plotter/</u>.

## 3. Results

Tybalt compressed tumors into a lower dimensional space, acting as a nonlinear dimensionality reduction algorithm. Tybalt learned which genes contributed to each feature, potentially capturing aberrant pathway activation and treatment vulnerabilities. Tybalt was unsupervised; therefore, it could learn both known and unknown biological patterns. In order to determine if the features captured biological signals, we characterized both sample- and gene-specific activation patterns.

### 3.1 Tumors were encoded in a lower dimensional space

The tumors were encoded from original gene expression vectors of 5,000 MAD genes into a lower dimensional vector of length 100. To determine if the sample encodings faithfully recapitulated large, tissue specific signals in the data, we visualized sample-specific Tybalt encoded features (z vector for each sample) by t-distributed stochastic neighbor embedding (t-SNE).^39^ We observed similar patterns for Tybalt encodings (Figure 2A) as compared to 0-1 normalized RNA-seq data (Figure 2B). Tybalt geometrically preserved well known relationships, including similarities between glioblastoma (GBM) and low grade glioma (LGG). Importantly, the recapitulation of tissue-specific signal was captured by non-redundant, highly heterogeneous features (Figure 2C). Based on the hierarchical clustering dendrogram, the features appeared to be capturing distinct signals. For instance, tumor versus normal and patient sex are large signals present in cancer gene expression, but they were distributed uniformly in the clustering solution indicating non-redundant feature activations.

### 3.2 Features represent biological signal

Our goal was to train and evaluate Tybalt on its ability to learn biological signals in the data and not to perform a comprehensive survey of learned features. Therefore, we investigated whether or not Tybalt could distinguish patient sex and patterns of metastatic activation. We determined that the model extracted patient sex robustly (Figure 3A). Feature encoding 82 nearly perfectly separated samples by sex. Furthermore, we identified a set of nodes that together identified skin cutaneous melanoma (SKCM) tumors of both primary and metastatic origin (Figure 3B).

The weights used to decode the hidden layer (z vector) back into a high-fidelity reconstruction of the input can capture important and consistent biological patterns embedded in the gene expression data.^7,8,32^ For instance, there were only 17 genes needed to identify patient sex (Figure 3C). These genes were mostly located on sex chromosomes. The two positive weight genes were X inactivation genes *XIST* and *TSIX*, while the negative weight genes were mostly Y chromosome genes such as *EIF1AY*, *UTY*, and *KDM5D*. This result served as a positive control that the unsupervised model was able to construct a feature that described a clearly biological source of variance in the data.

There were several genes contributing to the two encoded features that separated the SKCM tumors (Figure 3D). Several genes existed in the high weight tails of each distribution for feature encodings 53 and 66. We performed an ORA on the high weight genes. In general, several pathways were identified as overrepresented in the set as compared to random. The samples had intermediate to high levels of feature encoding 53, which did not correspond to any known GO term, potentially indicating an unknown but important biological process. The samples also had intermediate to high levels of encoding 66 which implicated GO terms related to cholesterol, ethanol, and lipid metabolism including regulation of intestinal cholesterol absorption (*adj. p* = 3.0*e*^0-2^), ethanol oxidation (*adj. p* = 4.0*e*^-02^), and lipid catabolic process (*adj. p* = 4.0*e*^-02^). SKCM samples had consistently high activation of both encoded features, which separated them from other tumors. Nevertheless, more research is required to determine how VAE features could be best interpreted in this context.

**Fig. 2.**
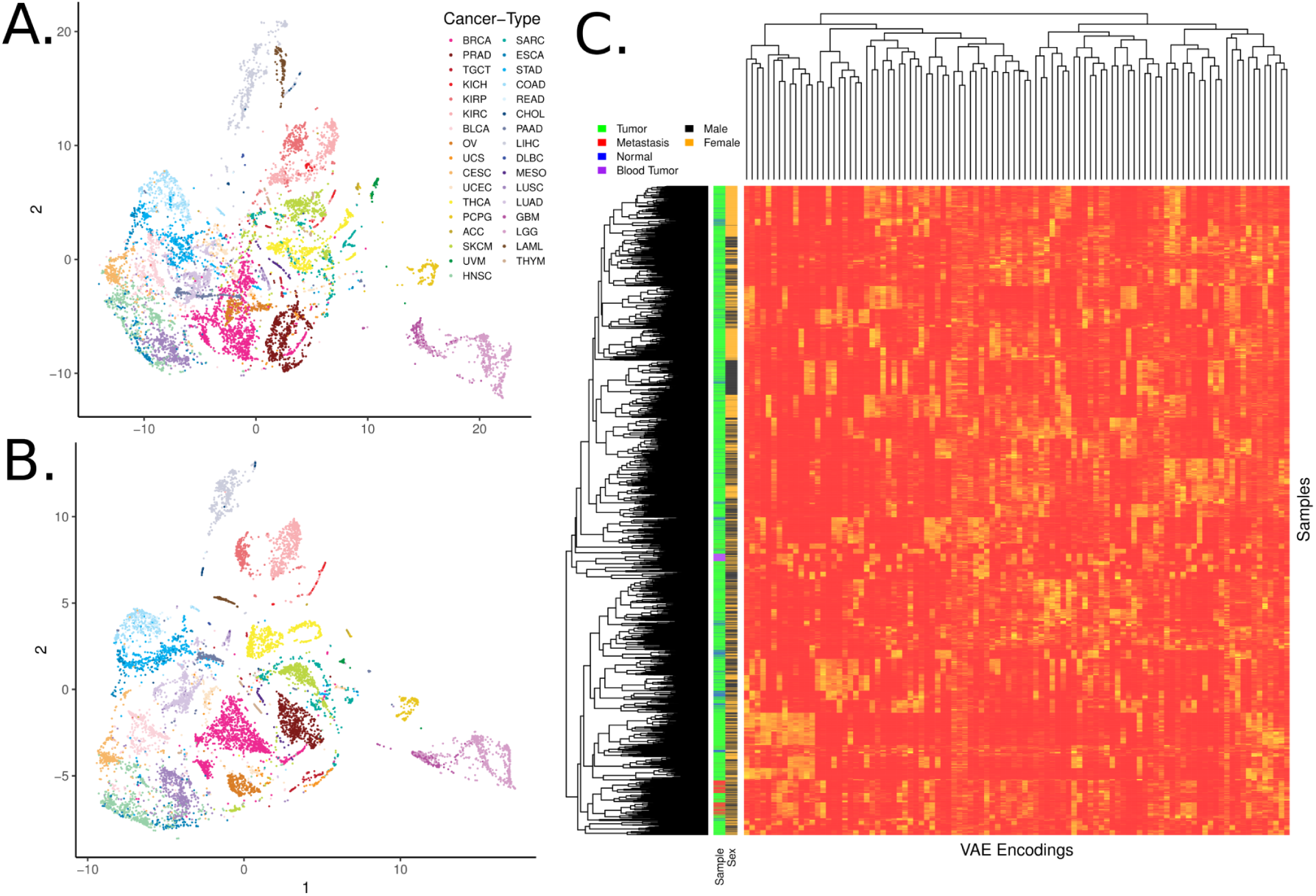
Samples encoded by a variational autoencoder retain biological signals. **(A)** t-distributed stochastic neighbor embedding (t-SNE) of TCGA pan-cancer tumors with Tybalt encoded features. **(B)** t-SNE of 0-1 normalized gene expression features. Tybalt retains similar signals as compared to uncompressed gene expression data. **(C)** Full Tybalt encoding features by TCGA pan-cancer sample heatmap. Given on the y axis are the patients sex and type of sample.

### 3.3 Interpolating the lower dimensional manifold of HGSC subtypes

We performed an experiment to test whether or not Tybalt learned manifold differences of distinct HGSC subtypes. Previously, several groups identified four HGSC subtypes using gene expression.^37,40,41^ However, the four HGSC subtypes were not consistently defined across populations; the data suggested the presence of three subtypes or fewer.^42^ The study observed that the immunoreactive/mesenchymal and differentiated/proliferative tumors consistently collapsed together when setting clustering algorithms to find 2 subtypes.^42^ This observation may suggest the presence of distinct gene expression programs existing on an activation spectrum driving differences in these subtypes. Therefore, we hypothesized that Tybalt would learn the manifold of gene expression spectra existing in differential proportions across these subtypes.

**Fig. 3.**
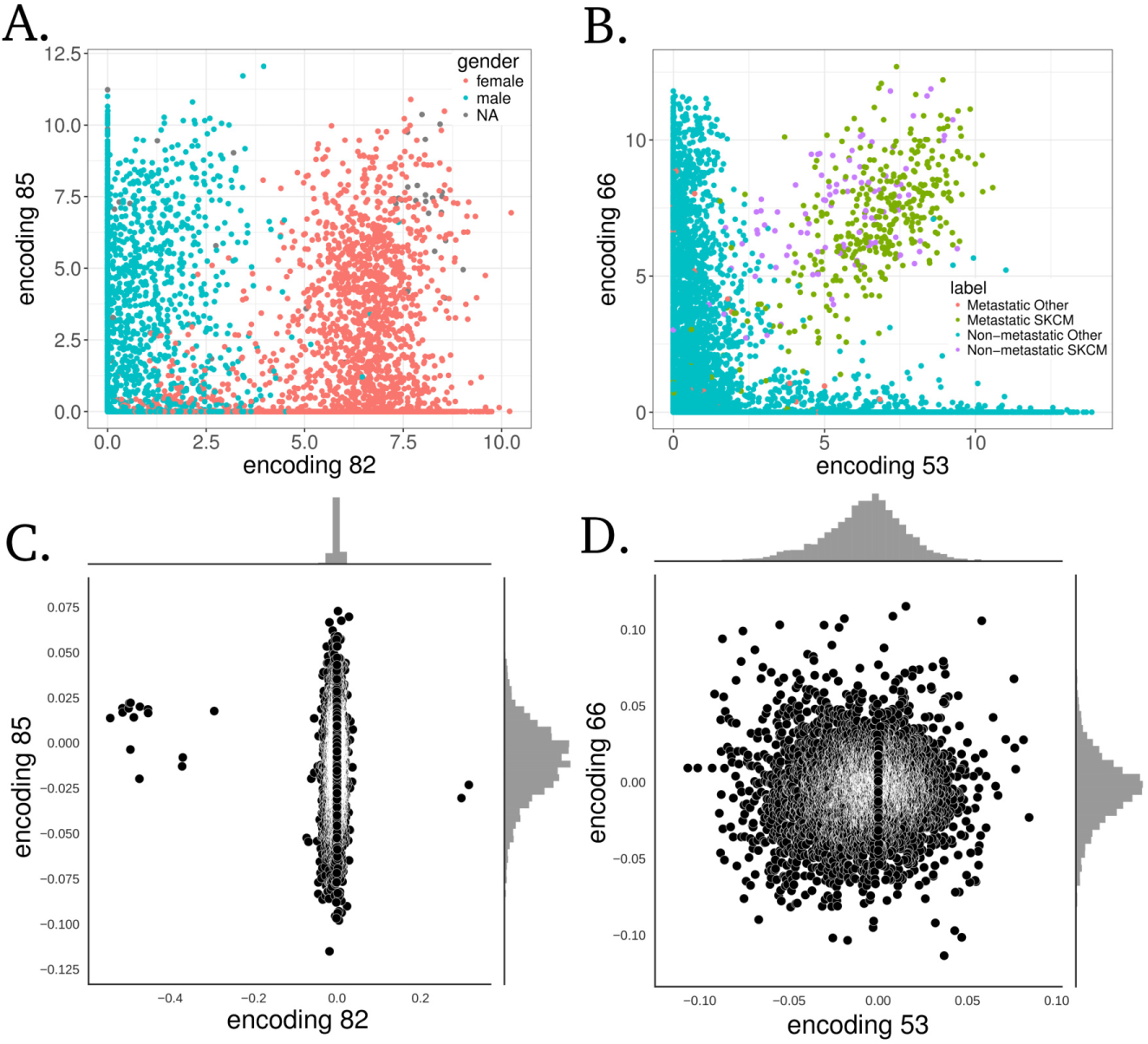
Specific examples of Tybalt features capturing biological signals. **(A)** Encoding 82 stratified patient sex. **(B)** Together, encodings 53 and 66 separated melanoma tumors. Distributions of gene coefficients contributing to each plot above for **(C)** patient sex and **(D)** melanoma. The gene coefficients consist of the Tybalt learned weights for each feature encoding.

The largest feature encoding difference between the mean HGSC mesenchymal and the mean immunoreactive subtype 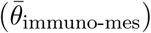 was encoding 87 (Figure 4A). Encoding 77 and encoding 56 (Figure 4B) also distinguished the mesenchymal and immunoreactive subtypes. The largest feature encoding differences between the mean proliferative and the mean differentiated subtype 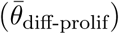 were contributed by encoding 79 (Figure 4C) and encoding 38 (Figure 4D). Interestingly, encoding 38 had high mean activation in both the immunoreactive and differentiated subtypes.

The mesenchymal subtype had the highest encoding 87 activation. Encoding 87 was associated with the expression of genes involved in collagen and extracellular matrix processes (Table 1), which has been previously observed to be an important marker of the mesenchymal subtype.^37,40^ Encoding 56 was associated with immune system responses (Table 1), and the immunoreactive subtype displayed the highest activation. Encoding 79 is mostly expressed in the proliferative subtype and has low activation in differentiated tumors. The high weight negative genes of encoding 79 were associated with glucuronidation processes (Table 1). The negative genes of encoding 38, which also distinguished differentiated from proliferative tumors but in the opposite direction, were also associated with glucuronidation. Previously, glucuronidation processes were observed to be associated with response to chemotherapy and survival in colon cancer patients.^43,44^ Our results indicate that differential activation of glucuronidation is a strong signal distinguishing HGSC subtypes. This observation may also help to explain increased survival in HGSC patients with differentiated tumors.^41^ Lastly, encoding 77 also separated immunoreactive from mesenchymal tumors and did not display any significant terms, which may indicate novel biology explaining undiscovered subtype differences.

**Fig. 4.**
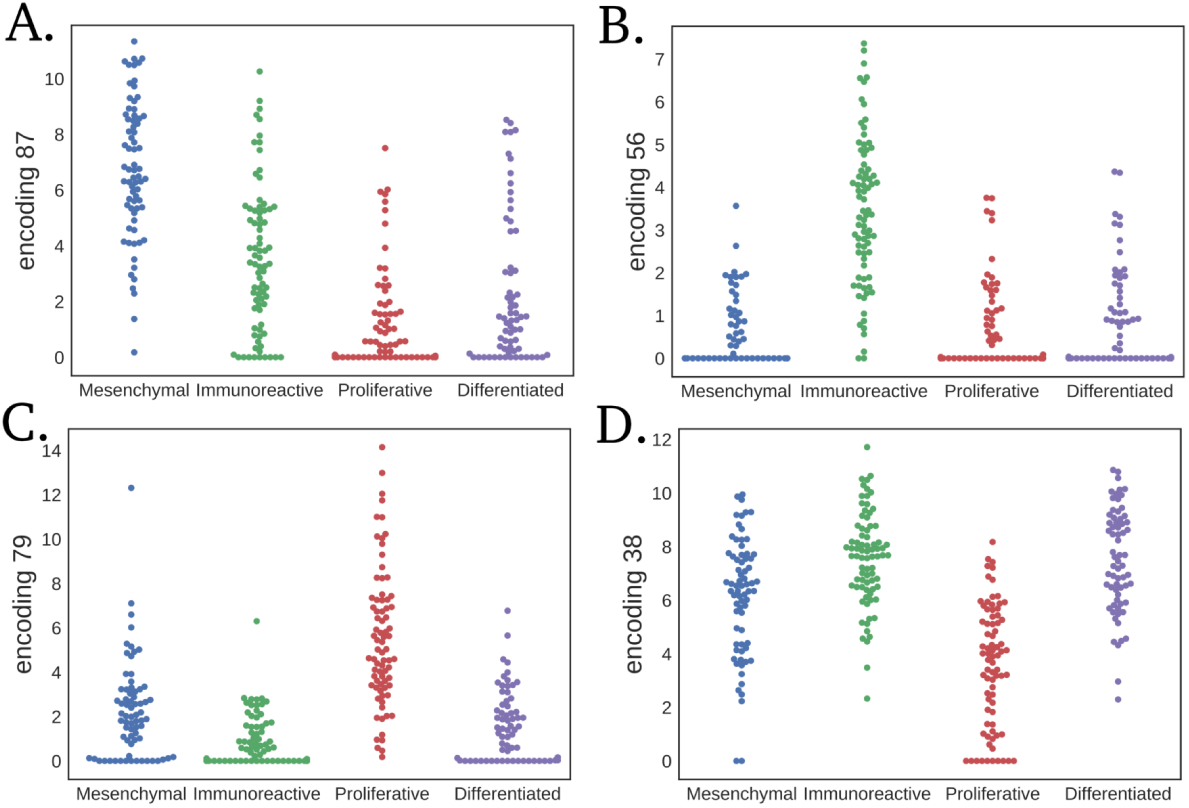
Largest mean differences in HGSC subtype vector subtraction for each subtype. Subtracting the mesenchymal subtype by the immunoreactive results in distribution differences in **(A)** feature encoding 87 and **(B)** encoding 56. Subtracting the proliferative subtype by the differentiated subtype results in differences between **(C)** feature encoding 79 and **(D)** encoding 38.

## 4. Conclusion

Tybalt is a promising model but still requires careful validation and more comprehensive evaluation. We observed that the encoded features recapitulated tissue specific patterns. We determined that the learned features were generally non-redundant and could disentangle large sources of variation in the data, including patient sex and SKCM. It is also likely that the features learn tissue specific patterns distinguishing other cancer-types (our shiny app enables full exploration of VAE features by cancer-type). While we identified specific features separating HGSC subtypes, there are likely several other features that describe other important biological differences across cancer-types including differentiation state and activation states of specific pathways. Interpretation of the decoding layer weights helped to identify the contribution of different genes and pathways promoting disparate biological patterns. However, interpretation by pathway analysis must be performed with caution as these analyses rely on incomplete pathway databases and may contain many false positive results.

**Table 1.** Summary of significantly overrepresented pathways separating HGSC subtypes

| Encoding | Tail | Subtype Enrichment | Pathway | Adj. p value |
| --- | --- | --- | --- | --- |
| 87 | + | Mesenchymal | Collagen Catabolic Process | $1.8e^{-09}$ |
| 87 | + | Mesenchymal | Extracellular Matrix Organization | $4.2e^{-06}$ |
| 87 | - | Immunoreactive | Urate Metabolic Process | $1.5e^{-02}$ |
| 56 | + | Immunoreactive | Immune Response | $1.3e^{-12}$ |
| 56 | + | Immunoreactive | Defense Response | $2.9e^{-12}$ |
| 56 | + | Immunoreactive | Regulation of Immune System Process | $8.0e^{-07}$ |
| 56 | - | Mesenchymal | *No significant pathways identified* |  |
| 79 | + | Proliferative | Chemical Synaptic Transmission | $9.1e^{-03}$ |
| 79 | - | Differentiated | Xenobiotic Glucuronidation | $2.1e^{-09}$ |
| 38 | + | Differentiated | *No significant pathways identified* |  |
| 38 | - | Proliferative | Xenobiotic Glucuronidation | $7.2e^{-06}$ |

VAEs provide similar benefits as autoencoders, but they also have the ability to learn a manifold with meaningful relationships between samples. This manifold could represent differing pathway activations, transitions between cancer states, or indicate particular tumors vulnerable to specific drugs. We performed initial testing to determine if we could traverse the underlying manifold by subtracting out cancer-type specific mean activations. While we identified several promising functional relationships existing in a spectrum of activation patterns, rigorous experimental testing would be required to draw strong conclusions about the biological implications. The specific subtype associations must be confirmed in independent datasets and the processes must be confirmed experimentally. It must also be assessed if Tybalt features learned from TCGA pan-cancer are generalizable to other, potentially more heterogeneous datasets. Further testing is required to confirm that Tybalt catalogued an interpretable manifold capable of interpolation between cancer states. In the future, we will develop higher capacity models and increased evaluation/interpretation efforts to catalog Tybalt encoded RNA-seq expression patterns present in specific cancer-types. This effort will lead to widespread stratification of expression patterns and enable accurate detection of samples who may benefit from specific targeted therapies.

## 5. Reproducibility

We provide all scripts to reproduce and to build upon this analysis under an open source license at https://github.com/greenelab/tybalt.^45^

## Acknowledgments

This work was supported by NIH grant T32 HG000046 (GPW) and GBMF 4552 from the Gordon and Betty Moore Foundation (CSG). We would like to thank Brett K. Beaulieu-Jones for helpful discussions and Jaclyn N. Taroni and David Nicholson for code review. We would also like to thank four anonymous reviewers for their insightful comments. This is a preprint of an article submitted for consideration in Pacific Symposium on Biocomputing (c)2018, World Scientific Publishing Co., http://psb.stanford.edu.

